# Deep learning-based prediction of wall shear stress throughout the cardiac cycle in middle cerebral artery bifurcation aneurysms

**DOI:** 10.64898/2026.10.12.758700

**Authors:** Bingyuan Bai, Xiajie Zhou, Kazunori Miyata, Tzu-Ching Shih

## Abstract

**Background and Objective:** Cardiac-cycle wall shear stress (WSS) is relevant to intracranial aneurysm hemodynamic assessment, but transient computational fluid dynamics (CFD) is costly. Existing surrogates use steady-flow priors or joint steady/transient training; here we evaluate the transfer of separately pretrained spatial parameters. This study develops the Steady-to-Transient Estimation of Pulsatile Wall Shear Stress (STEP) framework, a two-stage geometric learning approach for full-cycle WSS prediction.

**Methods:** Using the synthetic AneuG-Flow dataset, Stage 0 pretrained a geometric tokenizer, LaB-GATr backbone, and WSS decoder on 11,656 steady CFD cases. Stage 1 initialized these components from the pretrained model and fine-tuned them end to end using 449 transient training cases, with 72 validation and 57 primary test cases. Waveform embeddings multiplicatively conditioned geometric features across blocks at 80 cardiac-cycle time points. Gaussian curvature was included as an auxiliary geometry feature. Ablations evaluated steady initialization, temporal conditioning, and curvature, with rank-biserial effect sizes and Holm-adjusted *p* values. A fixed model was also evaluated on 231 additional cases, with registered-geometry overlap audited across stages.

**Results:** On the primary test set, STEP achieved a whole-cycle WSS vector mean absolute error (MAE) of 1.688 Pa and root-mean-square error (RMSE) of 2.877 Pa. Time-averaged WSS, oscillatory shear index, and relative residence time MAEs were 1.040 Pa, 0.00625, and 0.06734 Pa^−1^, respectively. Steady initialization reduced vector MAE by 27.93% relative to training the same transient architecture from scratch; the complete blockwise conditioning configuration reduced MAE by 8.04% relative to input-only additive conditioning (rank-biserial correlations 0.995 and 0.982, respectively; Holm-adjusted *p* < 0.001). Curvature yielded a 3.46% reduction without a significant paired effect (adjusted *p* = 0.0521). The 80-time-point WSS prediction required 4.169 ± 0.063 s. Additional-cohort MAE was 1.712 Pa, but 185/231 geometries matched Stage 0 training, versus 1/57 in the primary test set.

**Conclusions:** STEP demonstrated effective reuse of steady-pretrained spatial parameters for cardiac-cycle WSS prediction under the evaluated synthetic CFD protocol. The strongest controlled gains were associated with steady initialization and the complete temporal-conditioning configuration. The common inlet waveform and cross-stage geometry overlap limit generalization claims.

## I. Introduction

Intracranial aneurysms (IAs) are localized dilations of cerebral arteries associated with a risk of subarachnoid hemorrhage and substantial neurological morbidity.^1,2^ The management of unruptured IAs is informed by aneurysm size, anatomical location, and morphology, while local hemodynamic conditions provide additional biomechanical information.^3–5^ Wall shear stress (WSS), the tangential traction exerted by blood flow on the vessel wall, has been investigated in relation to aneurysm development, remodeling, and rupture status.^2,6^

WSS changes in magnitude and direction during pulsatile flow. Prediction throughout the cardiac cycle enables assessment of instantaneous surface loading and calculation of time-averaged WSS (TAWSS), oscillatory shear index (OSI), and relative residence time (RRT).^5,7,8^ Transient computational fluid dynamics (CFD) resolves these quantities but requires vascular geometry preparation, meshing, boundary-condition specification, and repeated numerical time stepping.^9,10^ The computational burden motivates a geometry-based surrogate capable of producing full-cycle WSS vectors without rerunning transient CFD for each evaluation.

Deep learning-based hemodynamic surrogates have used mesh learning, reduced-order representations, and geometry-conditioned networks to reduce the cost of flow-field estimation. MeshGraphNets established graph-based learning for physical simulations on unstructured meshes,^11^ while Rajhi et al. combined proper orthogonal decomposition with machine learning to reconstruct pulsatile hemodynamic fields in cerebral aneurysms.^12^ Gharleghi et al. predicted cardiac-cycle coronary WSS using vascular geometry and precomputed steady CFD information,^9^ and Suk et al. used steady-state hemodynamic priors to estimate pulsatile velocity and pressure.^13^ Geometry-driven models have also predicted peak-systolic velocity and WSS in synthetic cerebral aneurysms using point-cloud representations,^14^ time-averaged WSS in abdominal aortic aneurysms using multiview inputs,^15^ and coronary WSS in angiography-derived vessels using geometric deep learning.^16^ For vector-valued coronary WSS, Suk et al. developed an equivariant mesh network with waveform-conditioned full-cycle prediction.^17^

Recent studies have also investigated transient predictions and the use of prior geometric information in vascular surrogate modeling. Chan et al. compared CFD-supervised and physics-informed neural networks for steady and pulsatile flow in idealized vascular tubes,^18^ while Rygiel et al. studied geometry-based transient WSS estimation in abdominal aortic aneurysms under variable boundary conditions. ^19^ Lannelongue et al. developed a physics-constrained graph network for time-resolved intracranial aneurysm hemodynamics.^10^ Sheng et al. investigated full-cycle aneurysm WSS prediction on AneuG-Flow using graph-based architectures and joint steady/transient data augmentation, with waveform-derived features injected across transformer blocks.^20^ In a different pretraining setting, Suk et al. used self-supervised vascular shape representations to support coronary WSS-region assessment,^21^ whereas Hu et al. applied physics-guided implicit neural fields and progressive two-stage optimization to steady-state aneurysm velocity prediction. ^22^ These studies illustrate complementary approaches to spatial representation learning, temporal modeling, and pretraining.

An important question for full-cycle WSS prediction is whether spatial parameters learned from large-scale steady CFD supervision can provide an effective initialization for transient modeling. We investigate a sequential parameter-transfer strategy in which a geometry-to-WSS model is first pretrained using steady CFD data, and its learned spatial components are subsequently transferred to a waveform-conditioned transient model. Unlike the joint steady/transient augmentation investigated by Sheng et al., ^20^ steady training samples are not mixed into transient fine-tuning. Instead, the transferred parameters remain fully trainable, separating steady spatial initialization from temporal adaptation. This strategy reuses steady-supervised spatial knowledge without requiring case-specific steady CFD solutions at inference. A controlled comparison with the same transient architecture trained from scratch evaluates the contribution of steady initialization.

To implement this strategy, we develop the *Steady-to-Transient Estimation of Pulsatile Wall Shear Stress* (STEP) framework using the established Geometric Algebra Transformer (GATr) and its large-mesh variant LaB-GATr.^23,24^ Stage 0 learns geometric features and WSS decoding from steady supervision. Stage 1 transfers the pretrained geometric tokenizer, GATr backbone, and decoder, while a waveform encoder applies block-specific multiplicative scaling to geometric algebra features for time-resolved prediction. The complete temporal-conditioning configuration is evaluated against input-only additive conditioning under the same training protocol. Processed Gaussian curvature is incorporated as an auxiliary local shape descriptor alongside position and surface-normal information. Previous studies have reported associations between local surface curvature and WSS distributions. ^25,26^ In a CFD analysis of 76 patient-specific intracranial aneurysms, Dallavanzi et al. reported higher TAWSS and lower OSI in saddle-like surface regions than in spherical-like regions. ^27^ These findings provide a hemodynamic rationale for including explicit curvature information, whose incremental predictive contribution is evaluated separately.

We evaluate STEP on synthetic middle cerebral artery bifurcation aneurysms from the public AneuG-Flow resource.^28^ The study makes three contributions: (1) a two-stage steady-to-transient parameter-transfer framework, with a controlled scratch comparison to quantify the benefit of steady initialization; (2) a blockwise waveform-conditioning configuration for geometric algebra representations, evaluated against input-only additive conditioning; and (3) a comprehensive evaluation of complete 80-time-point WSS vectors, cycle-derived hemodynamic indices, high-WSS regions, computational efficiency, and registered-geometry overlap. The primary evaluation uses 57 transient test cases, with a separate 231-case cohort used for additional robustness assessment. Published source-study benchmarks provide descriptive performance context, while controlled component ablations establish the effects of the evaluated STEP configurations.

## II. Methods

STEP separates learning into two consecutive stages. Stage 0 trains a steady geometry-to-WSS spatial model; Stage 1 transfers and fine-tunes the spatial parameters while introducing waveform-derived conditioning for the complete 80-time-point WSS cycle (Fig. 1).

**Figure 1.**
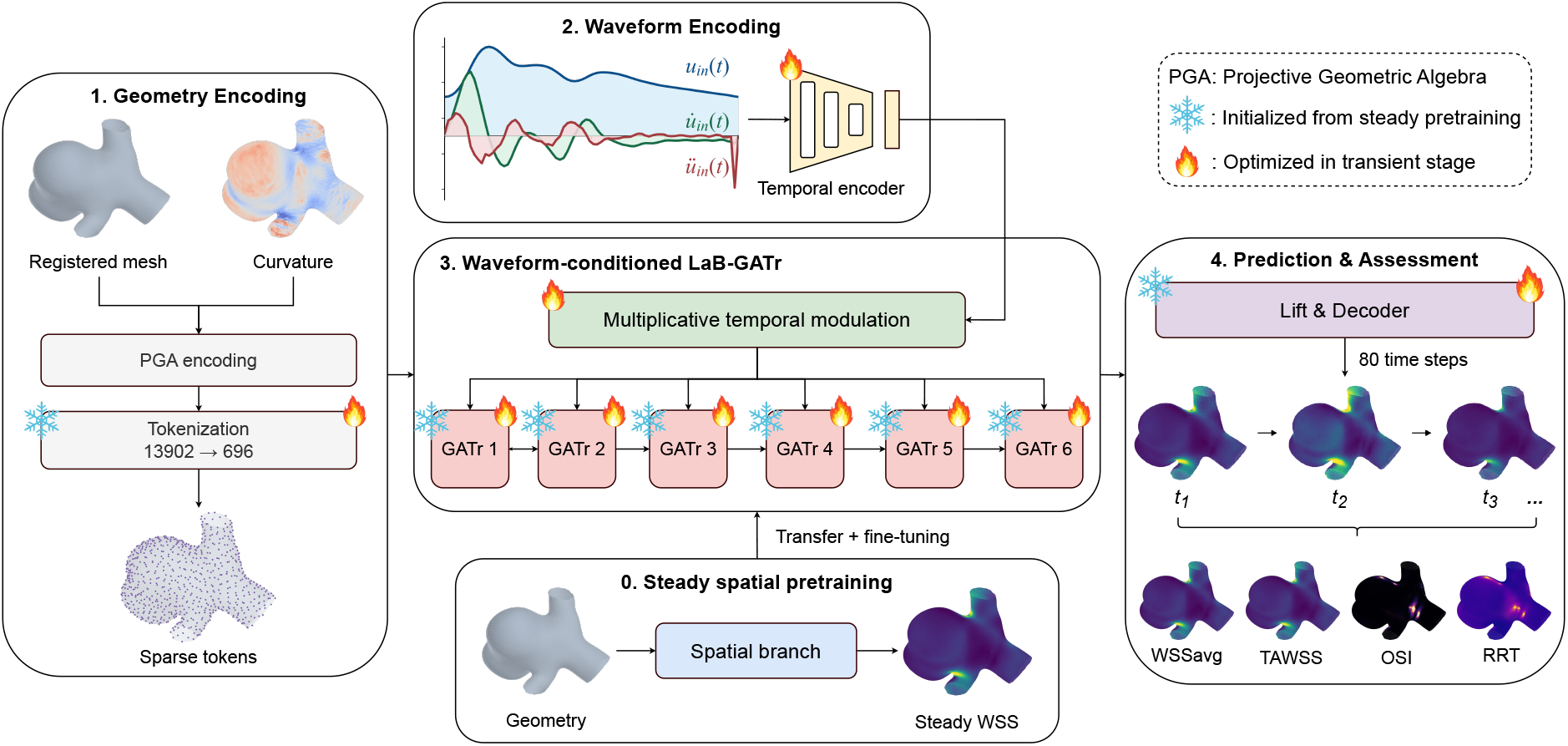
Overview of STEP, a two-stage steady-to-transient geometric learning framework. Stage 0 learns a spatial WSS representation from steady CFD data; Stage 1 initializes from the pretrained spatial branch and uses waveform-conditioned GATr blocks for cardiac-cycle WSS prediction and hemodynamic assessment.

### II.A. Dataset

The computational reference fields were obtained from the publicly released synthetic AneuG-Flow dataset. According to the source publication, CFD modeled blood as an incompressible non-Newtonian fluid using the Carreau–Yasuda model, with laminar flow, rigid no-slip walls, and a parabolic inlet profile of mean velocity 0.684 m s^*−*1^. A common pulsatile waveform modulated the inlet, while outlet flow rates followed a geometry-dependent modified Murray relation. Two cardiac cycles were simulated at a CFD time step of 0.001 s; 80 uniformly spaced time points were extracted from the second cycle. ^28^ Each registered transient surface contained 13,902 vertices with instantaneous three-dimensional WSS vectors, ***τ*** (**x**, *t*) (Pa). These CFD-derived fields served as model targets and were not patient-measured WSS or physiological ground truth; the common inlet waveform provided temporal conditioning. Source-patient identifiers were not available for the synthetically generated geometries, so patient-level independence could not be established from the released data. Case identifiers and registered-geometry matching were therefore audited as distinct, observable measures of data overlap.

The data comprised steady, transient, and transient extension components (Fig. 2(a)). The steady dataset contained 14,392 cases under the fixed seed-24 split: 11,656 training, 1,293 validation, and 1,443 test cases. Stage 0 optimization used only the 11,656 steady training cases, with the steady validation set used for model selection. Stage 1 used 578 transient cases. The author-aligned 57-case test split was retained unchanged, while the remaining 521 cases were divided into 449 training and 72 validation cases. ^20^ This primary transient test cohort is referred to as the 57-test set. The transient extension dataset contained 231 cases and is referred to as the 231-test set; no pulsatile fields from this cohort contributed to model development, and only the fixed validation-selected Stage 1 model was evaluated on it.

**Figure 2.**
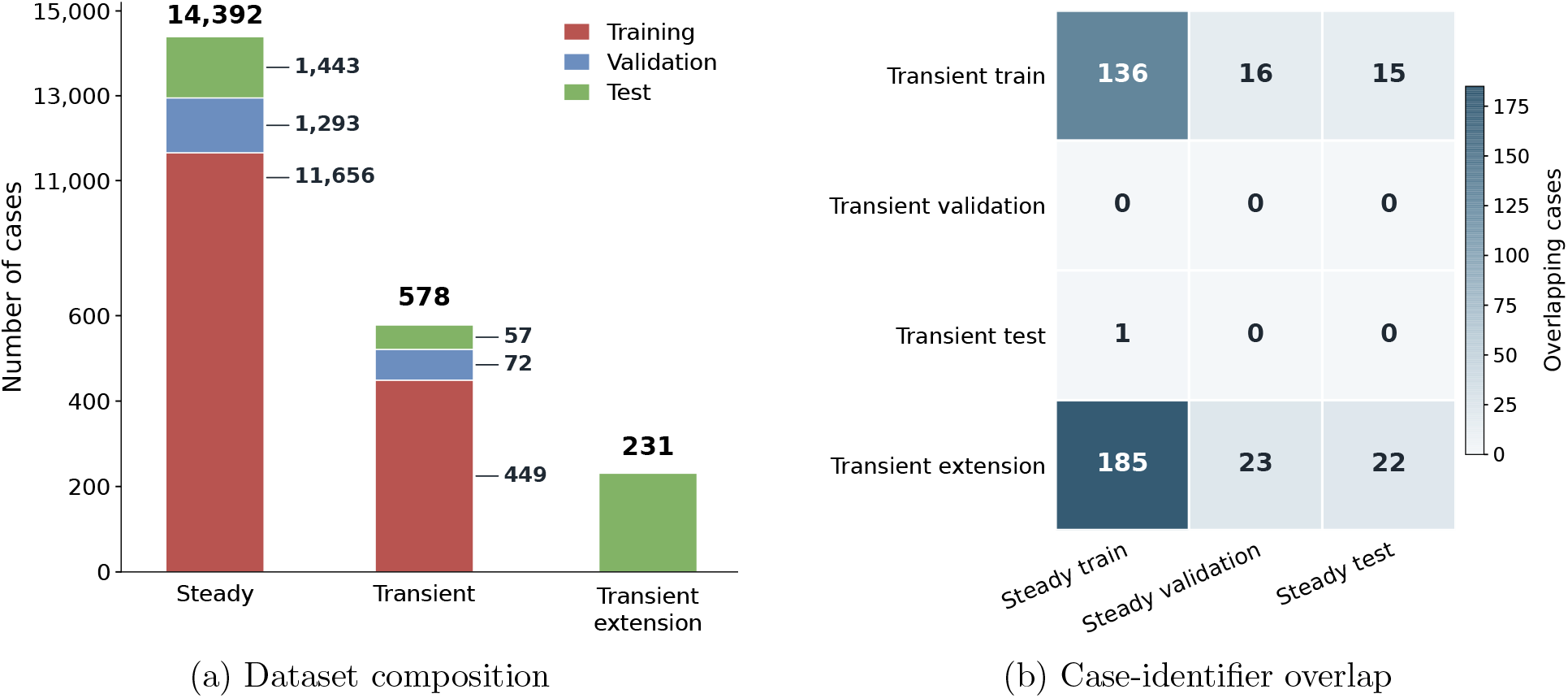
(a) Training, validation, and test composition of the steady, transient, and transient extension datasets. (b) Case-identifier overlap between the transient subsets and the corresponding steady-data partitions. Values indicate shared case identifiers only and do not establish registered-geometry identity. Registered-geometry overlap was audited independently and is reported in Sec. III.D..

Case identifiers and registered geometries were audited separately across the two-stage pipeline. Registered-geometry comparison used the complete 13,902-vertex coordinate arrays after reversing the stage-specific coordinate normalization and verifying the common 13,902-vertex, 27,604-face topology. A registered-geometry match required a maximum absolute coordinate difference no greater than 5 × 10^*−*5^ m and a coordinate root-mean-square error (RMSE) no greater than 2 × 10^*−*6^ m. Cross-stage comparisons included searches for registered-geometry matches across different case identifiers. Because different case identifiers were found to correspond to the same registered geometry, case-identifier overlap was not used as a substitute for registered-geometry overlap. Detailed overlap results and the corresponding sensitivity analysis are reported in Sec. III.D..

### II.B. Data preprocessing

The distributed coordinates used the source-dataset normalization **x**_norm_ = (**x**−*µ*_*r*_)*/*(*σ*_*r*_ +*ε*_*x*_), with *µ*_*r*_ = 2.9761402 × 10^*−*3^ m and *σ*_*r*_ = 9.4070588 × 10^*−*4^ m estimated from the complete 14,392-case steady dataset and applied identically to all Cartesian components. Following the released preprocessing, *ε*_*x*_ = 10^*−*6^ m for the steady data and 10^*−*5^ m for the transient data. Surface normals were normalized to unit length with tolerance 10^*−*5^.

Gaussian curvature *K* was computed on the normalized registered surfaces using PyVista 0.48.4 and the common triangular connectivity. Thus, *K* had inverse-square model-coordinate units and was not expressed in m^*−*2^. Open-boundary vertices were assigned zero curvature, and the remaining values were transformed as

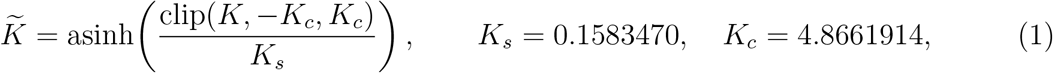

where *K*_*s*_ and *K*_*c*_ were the median and 99.5th percentile, respectively, of absolute interior curvature estimated exclusively from the 11,656 Stage 0 training cases and then fixed. 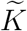 was used as an explicit scalar feature.

The normalized WSS components provided by the dataset were used directly as learning targets. Physical-unit evaluation used

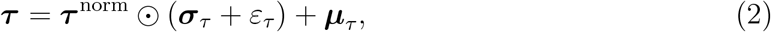

where ***µ***_*τ*_ = (−0.7324036, 1.1814325, 0.6199341) Pa, ***σ***_*τ*_ = (5.2252665, 4.8154626, 4.0063095) Pa, *ε*_*τ*_ = 10^*−*5^ Pa, and ⊙ denotes componentwise multiplication. These statistics were inherited from the complete steady dataset and were not re-estimated from transient data; no transient WSS targets contributed to their estimation.

For temporal conditioning, one 80,000-sample inlet-waveform cycle *u*_in_(*t*) was selected after removing the duplicate endpoint and resampled to *N* = 1000. With *T* = 0.8 s, Δ*t* = *T/N* = 8 × 10^*−*4^ s. Temporal derivatives were calculated by forward differences,

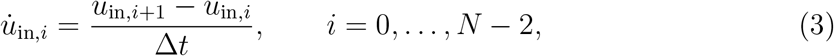

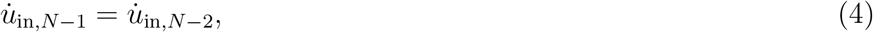

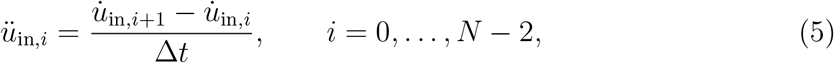

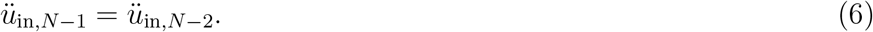

The channels 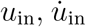, and *ü*_in_ had units of m s^*−*1^, m s^*−*2^, and m s^*−*3^, respectively. No additional z-score normalization was applied. The resulting 3 × 1000 temporal input and all other preprocessing parameters were shared across transient cases and applied without refitting.

### II.C. STEP framework: steady-to-transient learning

#### II.C.1. Steady spatial pretraining

For Stage 0 steady pretraining, each case was represented by

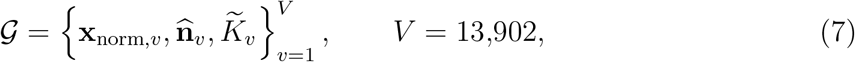

Where 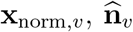, and 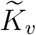 denote the normalized coordinate, unit surface normal, and processed Gaussian curvature at vertex *v*. The Stage 0 spatial branch comprised a deterministic projective geometric algebra (PGA) input encoder, a trainable geometric tokenizer, a six-block LaB-GATr backbone, deterministic full-resolution lifting with skip-feature concatenation, and a trainable WSS decoder. It was optimized against normalized steady CFD WSS using the loss in Sec. II.D.. Stage 1 initialized the tokenizer, GATr backbone, and decoder from the Stage 0 validation-selected checkpoint; all transferred parameters remained trainable, while the waveform encoder and temporal modulation gates were newly initialized.

#### II.C.2. Geometry encoding

For geometric encoding, the normalized coordinate was embedded as a PGA point and the surface normal with its position as an oriented plane:

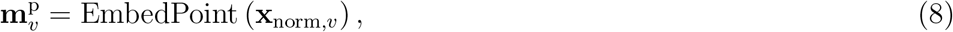

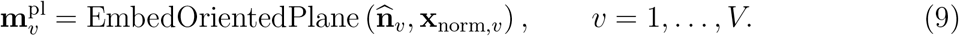

Both belong to **G**(3, 0, 1) and contain 16 Clifford-basis components.^23^ The full-resolution features were

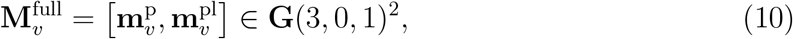

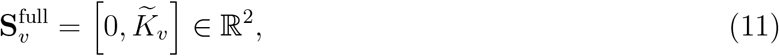

providing two multivector and two invariant scalar channels per vertex.

Deterministic farthest-point sampling (FPS) with random starts disabled selected 696 of the 13,902 vertices. Sampled features formed the query stream and the full-resolution features formed the key/value streams of the geometric tokenizer:

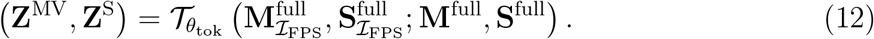

The tokenizer used a shared equivariant projection and four-head geometric cross-attention with normalization, residual updates, and a geometric multilayer perceptron (MLP). The 696 output tokens contained 32 multivector and two scalar channels; no class token or positional encoding was used. Full-resolution features were retained for the decoder skip pathway.

#### II.C.3. Waveform encoding

For waveform encoding, the 3 × 1000 waveform input 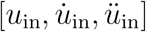 was processed by a three-layer one-dimensional convolutional encoder with channel sequence 3 → 32 → 64 → 32. Each convolution used kernel size 32, stride 2, and padding 16, with rectified linear unit (ReLU) activation; the sequence lengths were 501, 251, and 126. The encoded sequence was linearly interpolated to the *M* = 80 ordered cardiac-cycle time points and projected to a 32-dimensional temporal embedding:

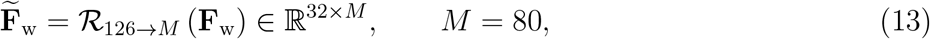

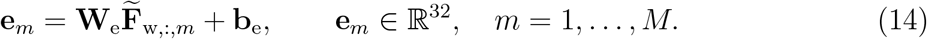

No separate time-index or positional embedding was introduced.

#### II.C.4. Waveform-conditioned transient GATr

For transient WSS prediction, the geometry tokens were projected to latent LaB-GATr features with 32 multivector and 128 scalar channels. The same geometry representation initialized every time point, and six four-head LaB-GATr blocks with dropout probability 0.05 processed the latent features.^24^ For each block *ℓ* = 1, …, 6, an independent linear gate mapped the time-point embedding to multivector- and scalar-channel scales:

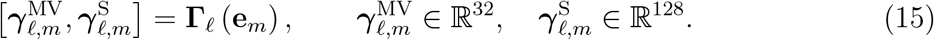

The scales were broadcast across the 696 tokens and applied immediately before the corresponding block:

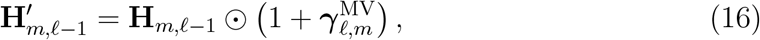

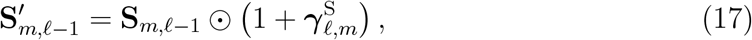

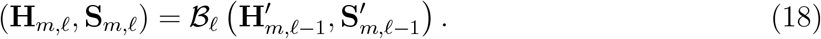

For each multivector channel, one invariant temporal scale was shared across all 16 Clifford-basis components; scalar features were scaled channelwise. The conditioning contained no additive temporal term. After the sixth block, an output projection returned each time-point-specific sparse representation to 32 multivector and two scalar channels for lifting.

#### II.C.5. Prediction and hemodynamic assessment

To recover full-resolution predictions, the 696 sparse tokens were lifted to the full 13,902-vertex surface by three-nearest-token inverse-distance interpolation. For sampled-token coordinate 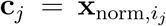 and the three-token neighborhood *N*_*v*_ of vertex *v*, the normalized weight was

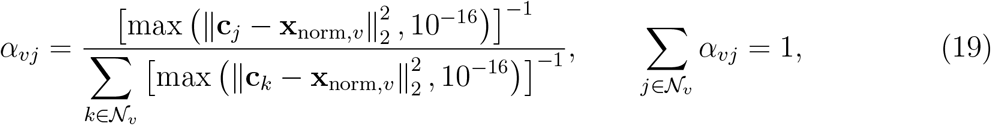

and the sparse multivector and scalar features were lifted as

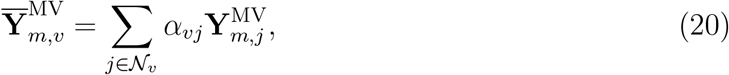

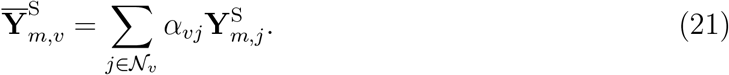

This interpolation contained no trainable parameters.

The lifted and full-resolution PGA features were concatenated and decoded by a geometric MLP with multivector widths 34 → 32 → 32 → 1, scalar-gated Gaussian error linear unit (GELU) activations, equivariant normalization at the second 32-channel stage, and a linear output projection. Translation extraction produced the normalized WSS prediction

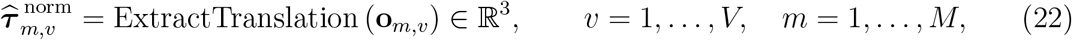

which was converted to physical units using Eq. (2).

The complete predicted cycle was post-processed as

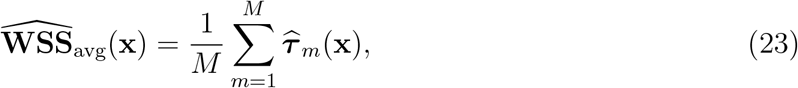

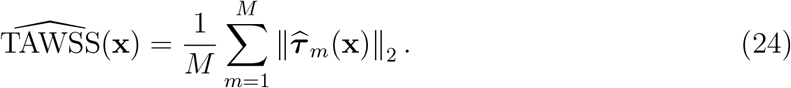

With *ε*_*h*_ = 10^*−*8^ Pa,

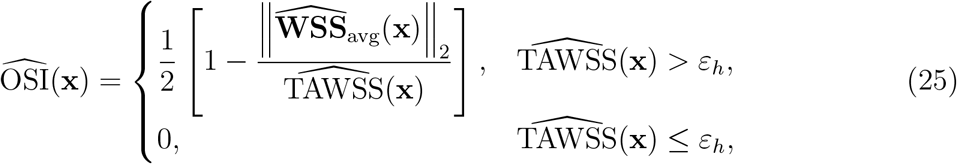

and

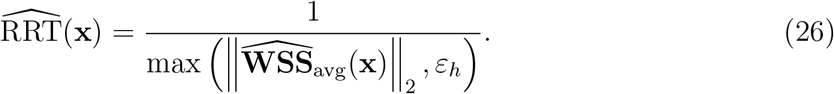

Reference WSSavg, TAWSS, OSI, and RRT were obtained from ***τ*** _*m*_(**x**) using the same definitions; reference OSI was additionally constrained to [0, 0.5] for compatibility with the frozen reference-field construction, whereas predicted OSI was not explicitly clipped. WSSavg and TAWSS are expressed in Pa, OSI is dimensionless, and RRT in Pa^*−*1^. All four quantities were derived from the complete 80-time-point sequence and were neither independent prediction heads nor training targets.

### II.D. Training configuration

Both stages were optimized on normalized three-component WSS using the componentwise L1 objective

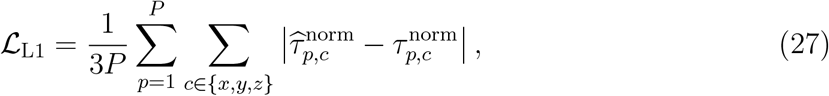

where *P* denotes the total number of vertices in the batch. No auxiliary supervision was imposed on WSSavg, TAWSS, OSI, RRT, or a specific temporal interval. Table 1 summarizes the optimization settings and graphics processing unit (GPU) hardware.

**Table 1:**
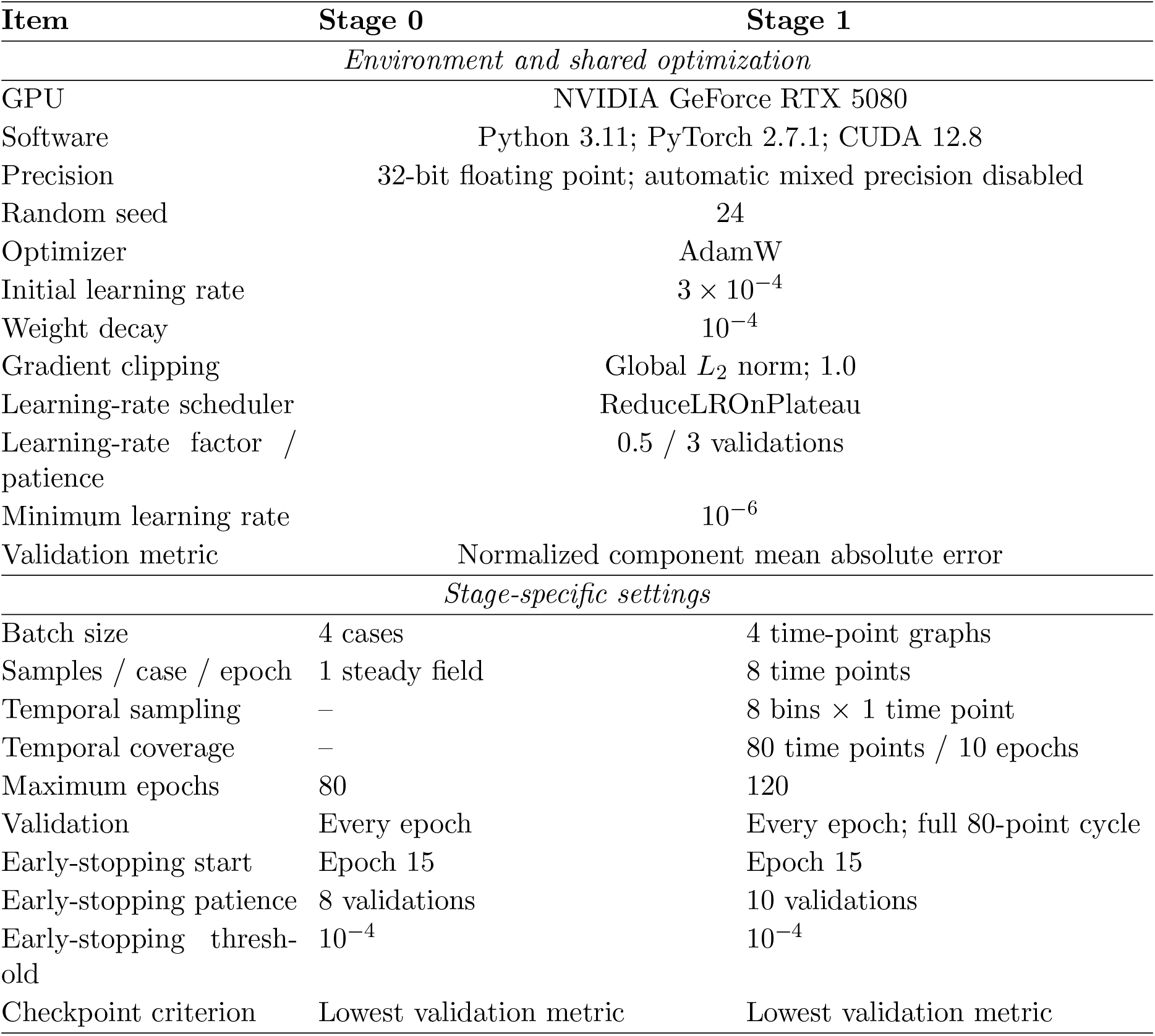
Optimization and stage-specific training configurations for the proposed framework: Stage 0 steady pretraining and Stage 1 transient training.

| Item | Stage 0 | Stage 1 |
| --- | --- | --- |
| *Environment and shared optimization* |  |  |
| GPU | NVIDIA GeForce RTX 5080 |  |
| Software | Python 3.11; PyTorch 2.7.1; CUDA 12.8 |  |
| Precision | 32-bit floating point; automatic mixed precision disabled |  |
| Random seed | 24 |  |
| Optimizer | AdamW |  |
| Initial learning rate | $3 \times 10^{-4}$ | |
| Weight decay | $10^{-4}$ | |
| Gradient clipping | Global $L_2$ norm; 1.0 | |
| Learning-rate scheduler | ReduceLROnPlateau |  |
| Learning-rate factor /<br>patience | 0.5 / 3 validations |  |
| Minimum learning rate | $10^{-6}$ | |
| Validation metric | Normalized component mean absolute error |  |
| *Stage-specific settings* |  |  |
| Batch size | 4 cases | 4 time-point graphs |
| Samples / case / epoch | 1 steady field | 8 time points |
| Temporal sampling | – | 8 bins × 1 time point |
| Temporal coverage | – | 80 time points / 10 epochs |
| Maximum epochs | 80 | 120 |
| Validation | Every epoch | Every epoch; full 80-point cycle |
| Early-stopping start | Epoch 15 | Epoch 15 |
| Early-stopping patience | 8 validations | 10 validations |
| Early-stopping thresh-<br>old | $10^{-4}$ | $10^{-4}$ |
| Checkpoint criterion | Lowest validation metric | Lowest validation metric |

For Stage 1, the cycle was divided into eight consecutive 10-point intervals, with one time point sampled from each interval per epoch. The sampling offset changed across epochs so that all 80 time points were covered once every 10 epochs. Validation used complete ordered 80-time-point cycles. The Stage 0 validation-selected checkpoint initialized Stage 1, and the Stage 1 validation-selected checkpoint was used for all transient test analyses. Neither 57-test nor 231-test participated in learning-rate scheduling, early stopping, or checkpoint selection.

### II.E. Ablation design

Three targeted component ablations assessed steady initialization, the complete blockwise temporal-conditioning configuration, and explicit curvature. Each comparator addressed only its corresponding design question; all unchanged data splits, optimization settings, validation procedures, and checkpoint criteria matched the proposed framework.

The scratch comparator retained the Stage 1 architecture but initialized its spatial parameters without loading the Stage 0 pretrained branch. The geometry-only comparator removed processed Gaussian curvature while retaining steady initialization and the complete temporal-conditioning architecture.

The temporal-modulation ablation retained curvature and steady initialization but replaced blockwise multiplicative modulation with a protocol-matched additive comparator. For time point *m*, the multivector features were unchanged, and the projected temporal embedding was added once to the initial scalar representation:

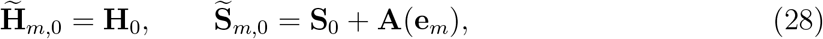

where **A** denotes the learned temporal projection. The six GATr blocks then operated without the per-block gates in Eq. (18).

A separate temporal-input ablation compared *u*_in_, 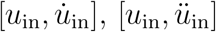, and the complete 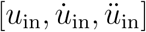 input. A temporal-sampling ablation evaluated 2, 4, 8, 16, and 32 sampled time points per case per Stage 1 epoch. Only the tested temporal input or sampling count was changed; the remaining Stage 1 protocol was retained. The final model used all three waveform channels and eight sampled time points per case per epoch. These two ablations were conducted only for descriptive analysis after the final model configuration had been fixed and did not inform model configuration or checkpoint selection.

### II.F. Surrogate assessment

#### II.F.1. Performance evaluation

Primary performance was assessed on 57-test over *M* = 80 time points. A sensitivity analysis repeated the primary evaluation after excluding the single case with a Stage 0-training registered-geometry match; the same validation-selected checkpoint, preprocessing, and metric definitions were retained for the remaining 56 cases. A separate numerical sensitivity analysis compared the WSS normalization statistics inherited from the complete steady dataset with statistics recalculated from the 11,656 Stage 0 training cases alone. Mean absolute error (MAE) measured average deviations, whereas RMSE emphasized larger vector errors. Vector magnitude MAE separated amplitude fidelity from directional agreement, characterized by cosine similarity. Let 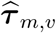 and ***τ*** _*m,v*_ denote the predicted and CFD-reference physical WSS vectors at time point *m* and registered vertex *v*, with magnitudes 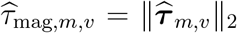 and *τ*_mag,*m,v*_ = ∥***τ*** _*m,v*_ ∥_2_.

For descriptive comparison with the source study,^20^ the released *rL*_2_ and 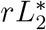 reductions were reproduced. Let *i* ∈ *J* index the ordered case–time-point–vertex evaluation stream. The first metric was

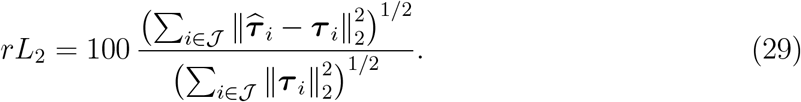

For ordered evaluation batches 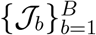 with *n*_*b*_ = |J_*b*_| and 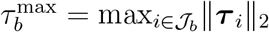,

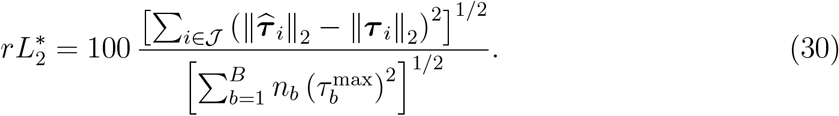

Both were evaluated in physical WSS units using batch size 16, the released evaluation order, and no shuffling, and were reduced globally over the evaluation stream without prior casewise reduction.

Instantaneous WSS accuracy was quantified for each case by

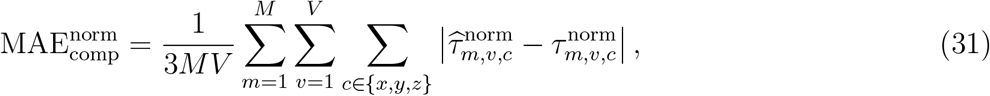

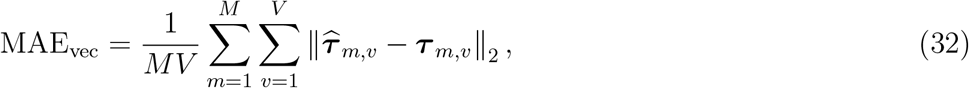

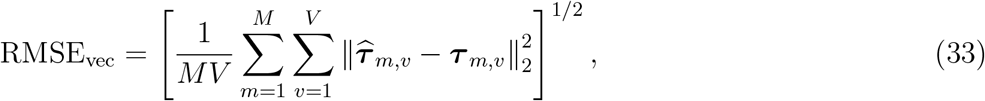

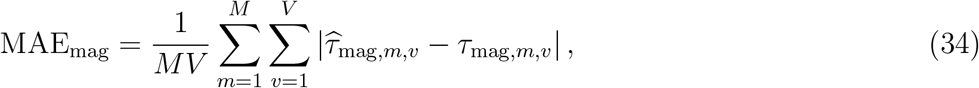

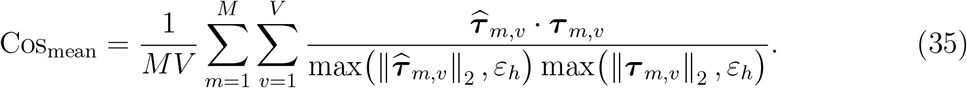

Normalized component MAE followed the Stage 1 training reduction; the remaining metrics used physical units. Except for *rL*_2_ and 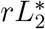, errors were reduced within each case before cohort-level summarization, with registered vertices weighted equally.

Cycle-derived accuracy used Eqs. (24)–(26):

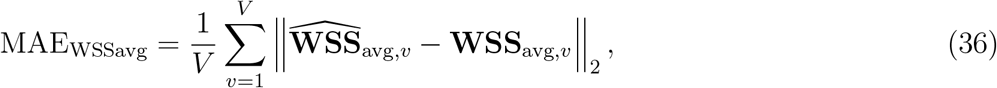

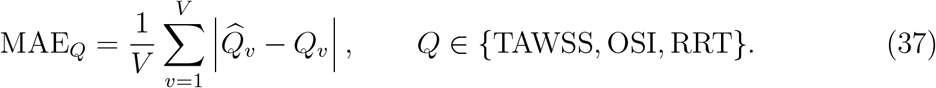

Time-resolved MAE_vec_ characterized temporal variation. Zero-based frame indices 12–18, corresponding to 0.12–0.18 s, defined the peak-inlet-velocity window, with maximum inlet velocity at frame index 12. Whole-cycle and peak-window errors used their corresponding time-point sets.

High-WSS fidelity was evaluated using CFD-reference 90th, 95th, and 99th percentile magnitude thresholds at each case and time point, defining top-10%, top-5%, and top-1% reference regions. The same reference threshold was applied to the prediction. Within the reference region, magnitude fidelity was quantified by the predicted-to-reference mean magnitude ratio and magnitude MAE: the ratio captured systematic over-or underestimation, whereas MAE captured absolute magnitude error. Spatial hotspot agreement was quantified by Dice similarity coefficient and intersection-over-union (IoU), using the same reference threshold for the prediction. Both metrics describe the same thresholded-region overlap and are retained for consistency with the hotspot visualization. Metrics were averaged across the relevant time points within each case and then summarized across 57-test, separately for the full cycle and frames 12–18.

Robustness was assessed by applying the fixed Stage 1 validation-selected model to 231-test without retraining, fine-tuning, or checkpoint reselection. The same preprocessing, complete-cycle prediction procedure, instantaneous and cycle-derived metrics, peak-window definition, and time-resolved analysis were retained. The 57-test versus 231-test comparison was descriptive.

#### II.F.2. Statistical analysis

The virtual aneurysm case was treated as the independent statistical unit. Continuous case-level quantities were summarized using the mean and standard deviation and the median and interquartile range as appropriate. Ninety-five percent confidence intervals (CIs) for case-level means and paired mean differences were estimated from 10,000 case-bootstrap resamples with seed 24. Time-resolved CIs used case-level cluster bootstrap resampling that preserved the complete within-case temporal sequence. Because model comparisons used a single training seed, these CIs quantify test-case sampling variability and do not quantify training-seed variability.

The three framework-component comparisons used whole-cycle MAE_vec_ on the same 57-test cases: full model versus geometry-only for curvature, full model versus scratch for steady initialization, and full model versus additive conditioning for temporal modulation. Paired differences were defined as comparator minus the full model, so positive values favor the full model. Two-sided Wilcoxon signed-rank tests were applied, with Holm correction across the three comparisons and adjusted *p <* 0.05 defining statistical significance. Paired rank-biserial correlation was reported as the effect size together with bootstrap 95% CIs for paired mean differences. Source-study comparison, temporal-input ablation, temporal-sampling ablation, and 57-test versus 231-test comparison were descriptive and received no additional hypothesis testing.

#### II.F.3. Computational efficiency

Computational cost was characterized for the final two-stage model using the hardware and software environment in Table 1. Stage-specific training time was obtained by summing recorded per-epoch durations. Peak allocated GPU memory was profiled separately for Stage 0 and Stage 1 using the corresponding final training configurations. Model size was reported as the total number of trainable parameters and the number of temporal-module parameters.

Inference runtime was measured for one complete 80-time-point cycle on a 13,902-vertex surface. Time points were processed in batches of four, requiring 20 forward passes per cycle. Three complete cycles were used for warm-up, followed by 10 timed repeats, with runtime summarized as the mean and sample standard deviation. Runtime included synchronous host-to-device transfer of pre-collated batches and model forward computation; disk input, geometry and waveform preprocessing, WSS denormalization, and WS-Savg/TAWSS/OSI/RRT post-processing were excluded. Peak allocated GPU memory was recorded over the timed inference repeats.

The AneuG-Flow source study reported approximately 3 min for mesh generation, 2 min for a steady CFD solve, and 10 h for each transient CFD solve. Each simulation was performed with ANSYS Fluent 2023R2 using 64 central processing unit (CPU) cores on a single AMD compute node.^28^

## III. Results

### III.A. Transient prediction

#### III.A.1. Primary performance

On the 57-case primary test set, STEP achieved a whole-cycle physical WSS vector MAE of 1.688 Pa and vector RMSE of 2.877 Pa. The peak-window MAE_vec_ over frame indices 12–18 was 3.795 Pa, with the largest cohort-mean time-resolved error occurring at frame index 13.

Cycle-derived errors on 57-test were 1.502 Pa for WSSavg vector MAE, 1.040 Pa for TAWSS MAE, 0.00625 for OSI MAE, and 0.06734 Pa^*−*1^ for RRT MAE. Representative spatial fields are shown in Fig. 3.

**Figure 3.**
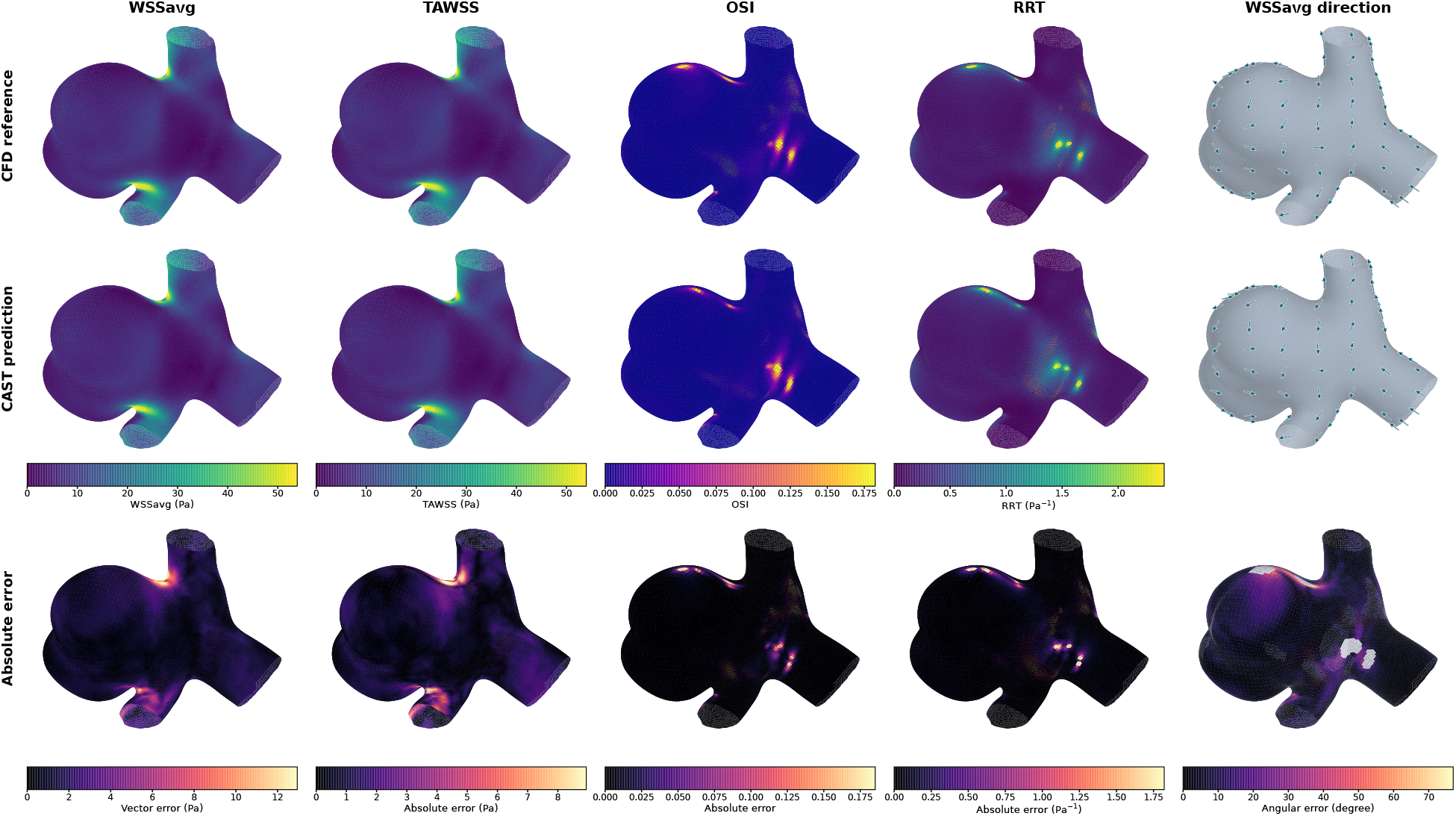
Computational-fluid-dynamics (CFD) reference, model-predicted, and error fields for the cycle-averaged WSS vector (WS-Savg), time-averaged WSS (TAWSS), oscillatory shear index (OSI), and relative residence time (RRT) in an example 57-test case. For WS-Savg, the reference and prediction panels show the magnitude of the cycle-averaged WSS vector, the error map shows the vector-difference norm, and the fifth column compares reference and predicted directions with their angular error. Vertices with reference WSSavg magnitude no greater than 2% of the case-specific 99.5th percentile of the reference WSSavg magnitude were excluded from the angular-error visualization. Display ranges were clipped at the 99.5th percentile.

#### III.A.2. Robustness

The fixed validation-selected model was applied to 231-test without retraining, fine-tuning, or additional checkpoint selection. Whole-cycle MAE_vec_ was 1.712 Pa (95% CI, 1.602–1.834), compared with 1.688 Pa on 57-test; RMSE_vec_ was 3.059 Pa (95% CI, 2.806–3.325), MAE_mag_ was 1.167 Pa (95% CI, 1.095–1.246), and mean cosine similarity was 0.9674. Peak-window MAE_vec_ was 3.836 Pa, compared with 3.795 Pa on 57-test.

Cycle-derived errors on 231-test were MAE_WSSavg_ = 1.517 Pa, MAE_TAWSS_ = 1.036 Pa (95% CI, 0.974–1.104), MAE_OSI_ = 0.00649 (95% CI, 0.00580–0.00730), and MAE_RRT_ = 0.06592 Pa^*−*1^. Case-level and time-resolved error profiles were similar across the two cohorts (Fig. 4); the maximum cohort-mean MAE_vec_ occurred at frame index 13 on 57-test (4.211 Pa) and frame index 14 on 231-test (4.258 Pa).

**Figure 4.**
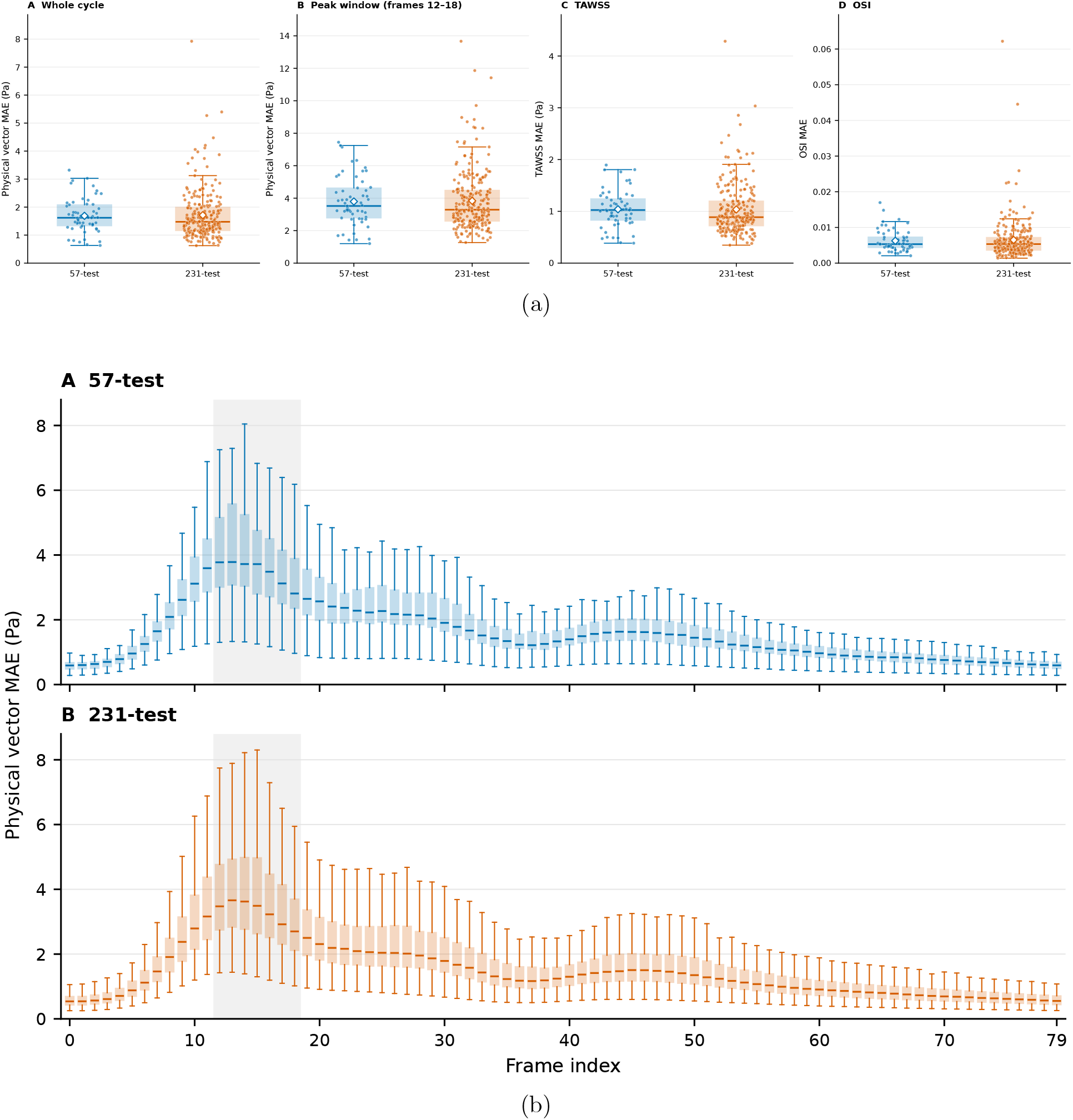
(a) Case-level distributions of whole-cycle MAE_vec_, peak-window MAE_vec_, MAE_TAWSS_, and MAE_OSI_ for 57-test and 231-test. Boxes indicate the median and interquartile range, whiskers extend to 1.5 times the interquartile range, points denote individual cases, and diamonds indicate cohort means. (b) Time-resolved MAE_vec_ across 80 uniformly spaced time points; the shaded interval denotes frame indices 12–18. MAE, mean absolute error; TAWSS, time-averaged wall shear stress; OSI, oscillatory shear index.

#### III.A.3. High-WSS fidelity

Across the complete cardiac cycle, predicted-to-reference magnitude ratios remained close to unity across all three percentile-defined regions, while spatial agreement decreased as the hotspot definition became more restrictive (Table 2). Performance decreased further during the peak-inlet-velocity window, with the top-1% region showing the largest magnitude error and the lowest spatial agreement. Representative hotspot localization is shown in Fig. 5.

**Table 2:** Case-level mean performance with 95% case-bootstrap confidence intervals on 57-test. High-WSS regions were defined by computational-fluid-dynamics (CFD) reference magnitude thresholds at the 90th, 95th, and 99th percentiles. WSS, wall shear stress; MAE, mean absolute error; Dice, Dice similarity coefficient; IoU, intersection-over-union.

| Time subset | Hotspot | Pred./reference<br>magnitude ratio | Magnitude MAE<br>(Pa) | Dice | IoU |
| --- | --- | --- | --- | --- | --- |
| Full cycle | Top 10% | 0.984 (0.972–0.997) | 2.473 (2.202–2.790) | 0.874 (0.861–0.886) | 0.781 (0.762–0.800) |
| Full cycle | Top 5% | 0.983 (0.969–0.998) | 3.021 (2.653–3.474) | 0.861 (0.847–0.874) | 0.761 (0.740–0.781) |
| Full cycle | Top 1% | 0.978 (0.958–0.998) | 4.312 (3.685–5.095) | 0.780 (0.748–0.809) | 0.657 (0.620–0.693) |
| Frames 12–18 | Top 10% | 0.967 (0.954–0.980) | 5.267 (4.730–5.837) | 0.818 (0.800–0.835) | 0.697 (0.673–0.721) |
| Frames 12–18 | Top 5% | 0.966 (0.950–0.982) | 6.309 (5.607–7.094) | 0.803 (0.783–0.822) | 0.678 (0.652–0.704) |
| Frames 12–18 | Top 1% | 0.954 (0.930–0.979) | 8.969 (7.793–10.368) | 0.700 (0.659–0.739) | 0.561 (0.518–0.603) |

**Figure 5.**
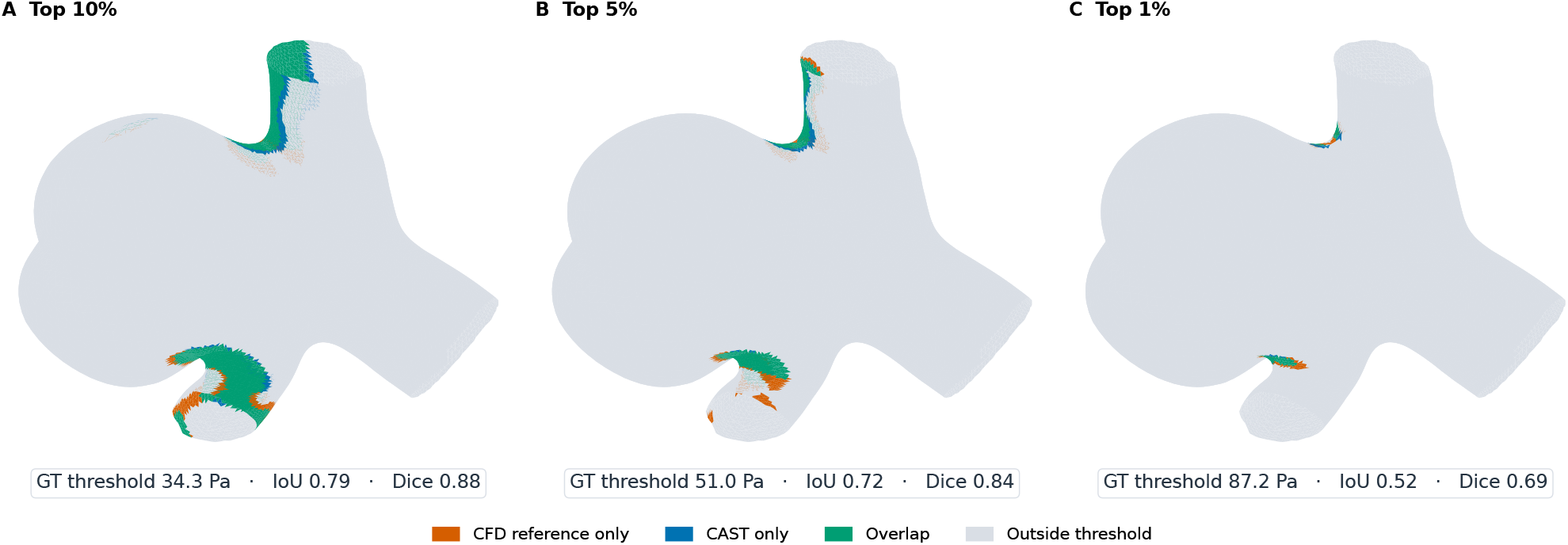
Computational-fluid-dynamics (CFD) reference and model-predicted top-10%, top-5%, and top-1% wall shear stress (WSS) hotspots in an example 57-test case at frame index 13. Threshold, Dice, and intersection-over-union (IoU) values in the panels correspond only to the illustrated case; cohort-level Dice and IoU results are reported in Table 2.

#### III.A.4. Source-study comparison

Table 3 lists the six transient WSS models reported in the AneuG-Flow source study together with the proposed model.^20^ Because the training protocols were unmatched and the metric provenance of the source-study values was not established from the released artifacts, no inferential comparison was performed.

**Table 3:** Author-reported transient wall shear stress (WSS) results for six source-study models and proposed-model results obtained using the released notebook reduction. C+N, coordinates and normals; PGA, projective geometric algebra; GHD, geodesic harmonic descriptors; CotLap, cotangent-Laplacian encoding; Curv., processed Gaussian curvature; n.r., not reported.

| Method | Input | Prediction | Temporal conditioning | Steady strategy | $rL_2$ (%) | $rL_2^*$ (%) |
| --- | --- | --- | --- | --- | --- | --- |
| U-Net | C+N | Framewise | Bottleneck | Augmentation | 29.51 | 3.20 |
| LaB-GATr | C+N | Framewise | n.r. | Augmentation | 36.78 | 4.55 |
| LaB-VATr | C+N | Framewise | n.r. | Augmentation | 28.04 | 3.40 |
| Sequence U-Net | C+N | Full-cycle | Cross-attention | Prior | 25.83 | 4.36 |
| Spectral Surrogate | GHD | Full-cycle | n.r. | Augmentation | 28.82 | 3.32 |
| Graph Transformer | C+N+GHD+CotLap | Framewise | Per-block | Augmentation | 26.98 | 2.84 |
| STEP (ours) | PGA(C+N)+Curv. | Framewise | Per-block multiplicative | Pretrain + fine-tune | 22.30 | 2.56 |

### III.B. Ablation studies

Figure 6(a) summarizes the three component ablations, with positive relative differences indicating lower error for the proposed model. Relative to the scratch, additive-conditioning, and geometry-only comparators, the proposed framework reduced physical WSS vector MAE by 27.93%, 8.04%, and 3.46%, respectively. Corresponding reductions were also observed for physical WSS vector RMSE, TAWSS MAE, and OSI MAE, with steady initialization producing the largest reduction among the three component comparisons across all four metrics.

**Figure 6.**
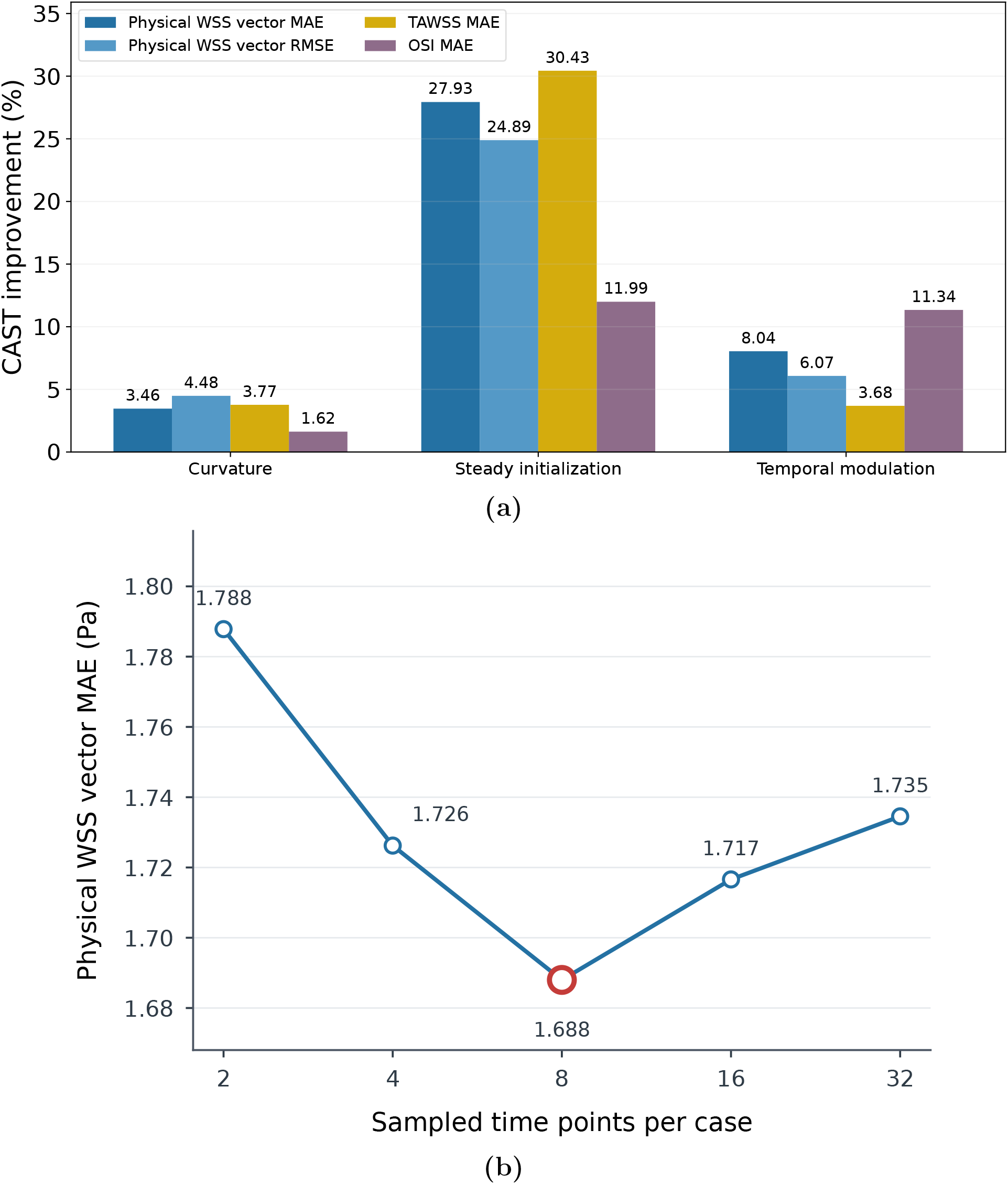
(a) Full-model relative error reduction compared with the corresponding curvature, steady-initialization, and temporal-modulation ablations; (b) physical wall shear stress (WSS) vector mean absolute error (MAE) across different numbers of sampled time points per case.

Across the temporal-input variants, the complete 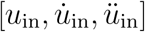 input yielded the lowest values across all four reported metrics, including a physical vector MAE of 1.688 Pa compared with 1.703 Pa for *u*_in_ alone (Table 4).

**Table 4:** Performance on 57-test for different combinations of waveform-derived temporal inputs. MAE, mean absolute error; RMSE, root-mean-square error; TAWSS, time-averaged wall shear stress; OSI, oscillatory shear index.

| Temporal input | MAE<sub>vec</sub><br>(Pa) | RMSE<sub>vec</sub><br>(Pa) | MAE<sub>TAWSS</sub><br>(Pa) | MAE<sub>OSI</sub><br>– |
| --- | --- | --- | --- | --- |
| $u_{\text{in}}$ | 1.703 | 2.900 | 1.052 | 0.00629 |
| $[u_{\text{in}}, \dot{u}_{\text{in}}]$ | 1.733 | 2.941 | 1.076 | 0.00632 |
| $[u_{\text{in}}, \ddot{u}_{\text{in}}]$ | 1.705 | 2.904 | 1.055 | 0.00633 |
| $[u_{\text{in}}, \dot{u}_{\text{in}}, \ddot{u}_{\text{in}}]$ | **1.688** | **2.877** | **1.040** | **0.00625** |

Physical WSS vector MAE varied nonmonotonically with temporal sampling and reached the lowest value among the evaluated settings with eight sampled time points per case (1.688 Pa; Fig. 6(b)).

### III.C. Statistical comparisons

#### III.C.1. Paired analysis of framework components

Steady initialization and the complete blockwise temporal-conditioning configuration significantly reduced case-level MAE_vec_ after Holm correction, whereas curvature did not reach statistical significance (Table 5).

**Table 5:** Paired case-level comparison of whole-cycle MAE_vec_ on 57-test. Mean paired differences were calculated as comparator minus the full model, such that positive values favor the full model. Confidence intervals were obtained from 10,000 case-bootstrap resamples, and *p* values were adjusted across the three comparisons using the Holm procedure. CI, confidence interval; MAE, mean absolute error.

| Component | Mean paired difference (95% CI)<br>(Pa) | Full model lower | Rank-biserial | Raw $p$ | Holm $p$ |
| --- | --- | --- | --- | --- | --- |
| Steady initialization | 0.654 (0.555–0.760) | 56/57 | 0.995 | $6.36 \times 10^{-11}$ | $1.91 \times 10^{-10}$ |
| Temporal conditioning | 0.148 (0.120–0.176) | 53/57 | 0.982 | $1.14 \times 10^{-10}$ | $2.28 \times 10^{-10}$ |
| Curvature | 0.060 (0.005–0.126) | 36/57 | 0.296 | 0.0521 | 0.0521 |

### III.D. Data-overlap audit and sensitivity

The coordinate-based audit refined the identifier-level overview in Fig. 2(b) by assessing registered-geometry overlap independently of case identifiers (Fig. 7). In 57-test, only 1/57 registered geometries matched Stage 0 steady training; comparison with the complete steady dataset identified one additional different-ID match located in the steady test partition. In the 231-test, 185/231 registered geometries matched Stage 0 steady training, so this cohort was retained only as an additional robustness cohort. The Stage 1 transient training and validation sets were audited separately, and neither had registered-geometry overlap with either transient test cohort. The two transient test cohorts also had no registered-geometry overlap with each other, and no duplicate registered geometries were identified within either cohort.

**Figure 7.**
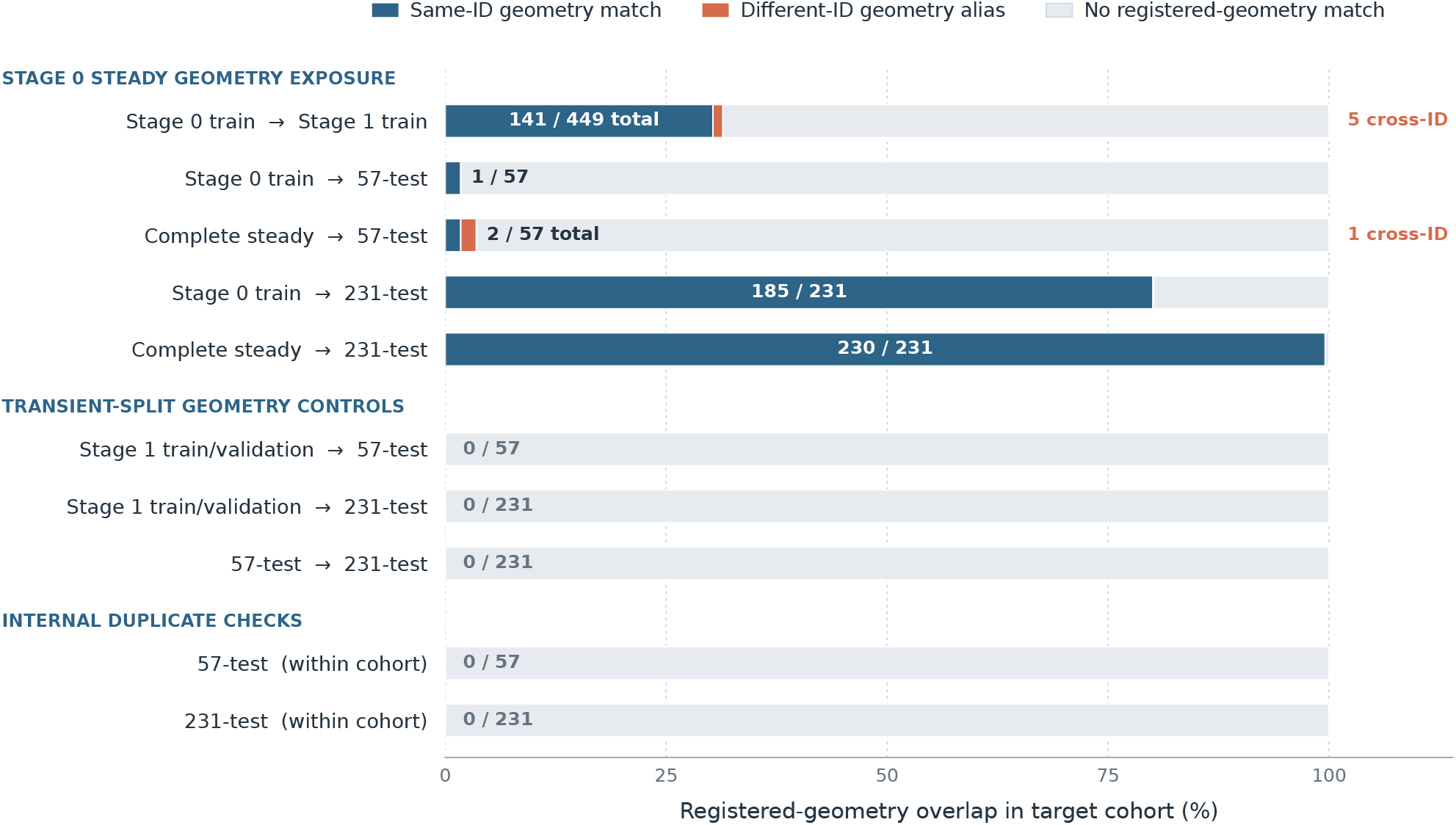
Registered-geometry overlap across the two-stage data pipeline. Bars distinguish same-ID geometry matches, different-ID geometry aliases, and cases without a registered-geometry match; fractions denote total matches relative to the target-cohort size.

To assess the effect of Stage 0 training geometry exposure on primary-test performance, we excluded the single matched 57-test case while retaining the same validation-selected checkpoint, preprocessing, and metric definitions. WSS normalization statistics derived from the complete steady dataset were also compared with statistics recalculated from the Stage 0 training set alone (Table 6).

**Table 6:** Sensitivity analysis after exclusion of the single Stage 0-training geometry-exposed case and comparison of WSS normalization statistics derived from the complete steady dataset and Stage 0 training set. The normalization difference denotes the maximum relative difference between the two sets of statistics. WSS, wall shear stress; MAE, mean absolute error; RMSE, root-mean-square error.

| Measure | Primary analysis | Sensitivity analysis | Difference |
| --- | --- | --- | --- |
| Cases | 57 | 56 | −1 |
| Stage 0-training geometry match | 1/57 | 0/56 | − |
| MAE<sub>vec</sub> (Pa) | 1.688 | 1.678 | −0.59% |
| RMSE<sub>vec</sub> (Pa) | 2.877 | 2.867 | −0.35% |
| WSS normalization statistics | Complete steady | Stage 0 training | 0.291% |

These sensitivity analyses did not alter the interpretation of primary-test performance.

### III.E. Computational efficiency

The proposed model achieved complete-cycle inference within seconds. Detailed training cost, model size, memory usage, and inference efficiency are summarized in Table 7.

**Table 7:** Training time and memory, model size, and inference efficiency of final two-stage model. Training times are sums of recorded per-epoch durations, and training-memory values were obtained from stage-specific profiling. Epochs are reported as the configured maximum, with the selected checkpoint epoch in parentheses. The transient CFD runtime is the author-reported value from the AneuG-Flow dataset study. ^28^ CFD, computational fluid dynamics; SD, standard deviation; GPU, graphics processing unit; CPU, central processing unit.

| Stage | Measure | Value |
| --- | --- | --- |
| Stage 0 | Maximum epochs (selected epoch) | 80 (77) |
|  | Training time | 23.96 h |
|  | Profiled peak allocated GPU memory | 6.292 GiB |
| Stage 1 | Maximum epochs (selected epoch) | 120 (49) |
|  | Training time | 6.60 h |
|  | Profiled peak allocated GPU memory | 6.345 GiB |
| Total training time |  | 30.56 h |
| Total trainable parameters |  | 2,726,067 |
| Spatial-backbone parameters |  | 2,559,059 (93.87%) |
| Temporal-module parameters |  | 167,008 (6.13%) |
| Complete-cycle inference time, mean ± SD | | 4.169 ± 0.063 s |
| Amortized runtime per time point |  | 52.1 ms |
| Inference peak allocated GPU memory |  | 1.29 GiB |
| Source-reported transient CFD solve time | | ∼10 h/case (64 CPU cores) |

## IV. Discussion

STEP demonstrated effective transfer of steady-trained spatial parameters for cardiac-cycle WSS prediction under the evaluated synthetic CFD protocol. Steady initialization produced the largest improvement among the controlled component comparisons, followed by the complete blockwise temporal-conditioning configuration; both effects remained significant after Holm correction. Auxiliary curvature did not yield a statistically significant case-level improvement. These findings support the implemented two-stage learning strategy without claiming that staged learning or waveform conditioning was introduced for the first time.

Previous hemodynamic surrogates have used computed steady fields as inputs or priors.^9,13^ Sheng et al. investigated joint steady/transient augmentation for aneurysm WSS prediction.^20^ STEP instead transfers pretrained tokenizer, LaB-GATr backbone, and WSS decoder parameters into Stage 1, where they remain fully trainable. The controlled scratch comparison establishes the benefit of this initialization within the same transient architecture and training protocol. It does not establish superiority over joint augmentation, which was not evaluated under a matched protocol; the source-study results in Table 3 therefore provide descriptive benchmark context.

Waveform-based per-block injection and higher-order waveform derivatives have been investigated previously.^20^ STEP uses block-dependent multiplicative scales shared across the Clifford components of each multivector channel, with separate scaling of scalar channels. The full configuration reduced whole-cycle vector MAE by 8.04% relative to input-only additive conditioning, but this comparison also changes the injection depth and does not isolate multiplication as the cause. The complete 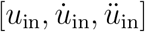 input achieved the lowest error among the tested waveform representations, although individual derivatives did not consistently improve on the inlet waveform alone (Table 4). Eight sampled time points per case per epoch performed best among the tested sampling settings (Fig. 6(b)); neither observation establishes a new temporal-conditioning principle.

Gaussian curvature was an auxiliary descriptor of second-order surface geometry beyond the positions and normals encoded through PGA. Its inclusion reduced aggregate vector MAE by 3.46% compared with the geometry-only model, but the Holm-adjusted paired comparison was not significant (*p* = 0.0521; Table 5). The positive bootstrap confidence interval for the mean paired difference and nonsignificant Wilcoxon result address different aspects of the paired error distribution; our inferential conclusion follows the corrected Wilcoxon test.

The endpoint-level evaluation illustrates both the utility and limitations of predicting complete WSS cycles. WSSavg, TAWSS, OSI, and RRT were derived from the predicted sequence without separate task-specific heads. Errors increased near peak inlet velocity, and both localization and magnitude fidelity weakened in restrictive high-WSS regions, particularly during frames 12–18 (Sec. III.A.3.; Table 2); whole-surface mean error therefore does not capture all localized extremes. Model inference for 80 time points on 13,902 surface vertices required 4.169 ± 0.063 s, with temporal modules accounting for 6.13% of the trainable parameters. Stage 0 training is an upfront cost, not a per-case inference cost. A CFD acceleration factor is not reported because the surrogate and CFD runtimes were obtained on unmatched hardware and workloads.

The fixed model produced similar error summaries on the additional 231 transient cases without retraining or checkpoint reselection, but this cohort cannot establish fully unseen-geometry generalization across both training stages: 185/231 registered geometries matched Stage 0 training. By comparison, 1/57 primary-test geometries matched Stage 0 training, and excluding that case did not materially change aggregate performance (Sec. III.D.). No patient-level identifiers were available, and the registered-geometry audit cannot establish broader anatomical independence.

Several limitations qualify these findings. **(1)** Reference WSS fields were generated by synthetic CFD under rigid-wall, rheological, inlet, and outlet assumptions; agreement with them is not evidence of physiological or clinical accuracy. **(2)** A common inlet waveform was used for all transient cases, leaving generalization to different waveform shapes, amplitudes, heart rates, and outlet conditions untested. **(3)** The registered-geometry audit does not establish patient-level independence or fully unseen anatomy. **(4)** Evaluation was restricted to synthetic middle cerebral artery bifurcation aneurysms; other locations and imaging or segmentation variability were not tested. **(5)** Patient outcomes, predictive uncertainty, and clinical decision utility were not assessed. **(6)** The temporal-conditioning comparator differs in both fusion operation and injection depth, so the isolated effect of multiplication remains unestablished. Further work should assess heterogeneous boundary conditions, improve high-WSS fidelity during peak flow, and validate broader geometry cohorts and clinical endpoints before clinical application.

## V. Conclusion

STEP combines steady WSS pretraining with waveform-conditioned, end-to-end transient fine-tuning for full-cycle WSS prediction in synthetic middle cerebral artery bifurcation aneurysms. On the 57-case primary test set, steady initialization reduced vector MAE by 27.93% relative to scratch training, and the complete blockwise temporal-conditioning configuration reduced MAE by 8.04% relative to input-only additive conditioning. Curvature provided no statistically significant paired improvement. Model inference for 80 time points required approximately 4.17 s on the evaluated GPU. The additional 231-case results provide robustness context, but Stage 0 geometry overlap and the common inlet waveform limit generalization claims. These findings establish the effectiveness of the implemented two-stage strategy under the evaluated synthetic CFD protocol.

## Data Availability Statement

The AneuG-Flow data analyzed in this study are publicly available from the source dataset described in Ref.^28^. The code developed for this study is available from the corresponding author upon reasonable request.

## Author Contributions

Bingyuan Bai: Methodology, Software, Formal analysis, Investigation, Visualization, Writing–original draft. Xiajie Zhou: Data curation, Investigation. Kazunori Miyata: Project administration, Resources. Tzu-Ching Shih: Conceptualization, Supervision, Resources, Funding acquisition, Writing–review and editing. All authors critically reviewed the manuscript for important intellectual content and approved the final version.

## Acknowledgments

This work was supported by the National Science and Technology Council (NSTC), Taiwan, under Grant Nos. 115-2314-B-039-035 (Multimodal Fluid–Structure Interaction Coupled with the TESLA Framework for Personalized Prediction of Intracranial Aneurysm Rupture Risk), and 112-2221-E-039-003-MY2.

## Declaration of Generative AI and AI-assisted Technologies in the Writing Process

During the preparation of this manuscript, the authors used OpenAI ChatGPT for English-language grammar checking and sentence-level polishing. All AI-assisted edits were reviewed and verified by the authors, who take full responsibility for the final manuscript.

